# BioKC: a collaborative platform for systems biology model curation and annotation

**DOI:** 10.1101/2020.10.01.322438

**Authors:** Carlos Vega, Valentin Grouès, Marek Ostaszewski, Reinhard Schneider, Venkata Satagopam

## Abstract

Curation of biomedical knowledge into standardised and inter-operable systems biology models is essential for studying complex biological processes. However, systems-level curation is a laborious manual process, especially when facing ever increasing growth of domain literature. Currently, these systems-level curation efforts concentrate around dedicated pathway databases, with a limited input from the research community. The demand for systems biology knowledge increases with new findings demonstrating elaborate relationships between multiple molecules, pathways and cells. This new challenge calls for novel collaborative tools and platforms allowing to improve the quality and the output of the curation process. In particular, in the current systems biology environment, curation tools lack reviewing features and are not well suited for an open, community-based curation workflows. An important concern is the complexity of the curation process and the limitations of the tools supporting it. Currently, systems-level curation combines model-building with diagram layout design. However, diagram editing tools offer limited annotation features. On the other hand, text-oriented tools have insufficient capabilities representing and annotating relationships between biological entities. Separating model curation and annotation from diagram editing enables iterative and distributed building of annotated models. Here, we present *BioKC* (<u>Bio</u>logical <u>K</u>nowledge <u>C</u>uration), a web-based collaborative platform for the curation and annotation of biomedical knowledge following the standard data model from Systems Biology Markup Language (SBML).

## Introduction

Since the beginning of computational systems biology during the analogue computer era [1, 2], researchers aim to formalise biological processes into computational models for their analysis and simulations. The process of curation in systems biology is time-consuming, requires domain knowledge to explore, organise, and encode the abundance of information available in the literature, and often involves several experts in the process. In recent years the demand for systems biology knowledge increases (see Fig. 1), following scientific advances and improved methods for analysis of complex systems and data sets.

**Fig. 1.**
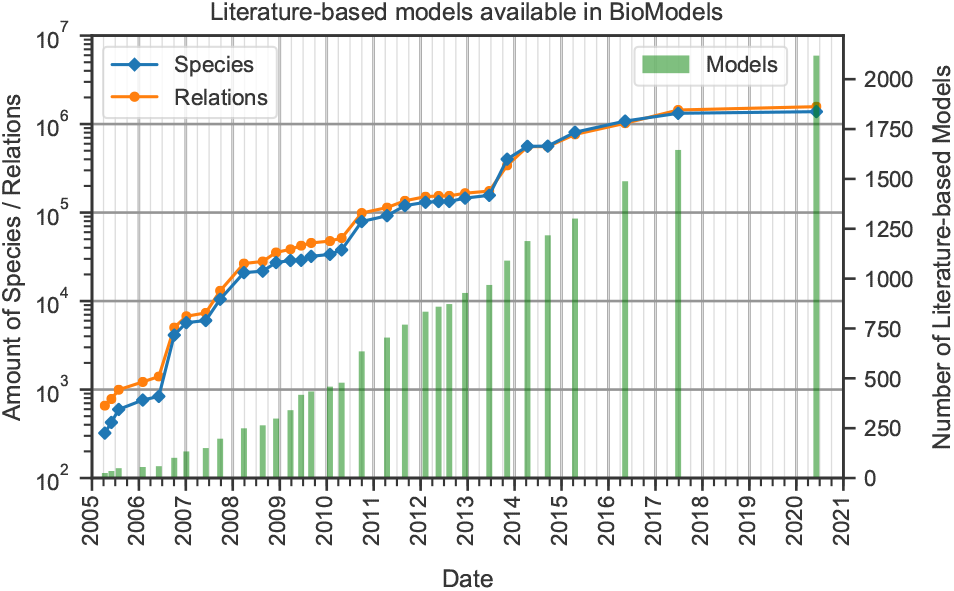
Evolution of literature-based models available in BioModels repository [9].

However, few tools feature collaborative web functionalities for systematic and quality controlled model curation. In effect, current curation efforts focus around a limited number of groups associated with pathway and interaction databases. Moreover, systems biology tools like diagram editors or publication annotation tools are not the best suited for high quality curation. Systems biology modelling tools have different strengths and capabilities [3], ranging from diagram editors like CellDesigner [4] or Newt [5] to tools relying on text files structured following a given standard, like Cytoscape [6] or COPASI [7]. Despite their different nature and purpose, most of these tools are interoperable, thanks to representation formats such as the Systems Biology Markup Language (SBML) [8]. Nevertheless, when it comes to tasks related to model building and annotation, they lack in terms of high quality curation workflows. On one hand, tools with well-developed graphical user interfaces have limited capabilities to support in-depth curation, e.g. handling supporting sentences from scientific articles or versioning the model. On the other hand, tools relying on text interfaces and representation formats allow for detailed annotation, but offer limited interfaces to support the work of curators (reviewed in Section <u>*Related work*</u>).

### KEY COMPONENTS OF A CURATION WORKFLOW

Curators of systems biology models: (i) extract bio-entities, relationships and annotations from the literature to build a model; (ii) craft a consistent textual or graphical representation of such model; and (iii) review and parameterise the model. Because of a number of available tools and modelling approaches, a biological process may be encoded in various system biology formats, and then depicted in different ways. Developing a diagrammatic representation of the model requires additional effort and dedicated tools. Diagram editors, supporting this task, aggregate model definition, layout, and annotations in a monolithic block. This in turn hinders reusability, extension and management of such models [10], in particular of the annotations and provenance tracking of the literature used to establish the model. Moreover, a particular biological process may need different graphical representations depending on the context. For an existing diagram, it requires copying the model annotations and modifying the layout. In such a scenario, annotation management and change tracking becomes tedious and error-prone.

Thus, key components of the curation workflow should consider: (i) a model, representing biomedical elements and their interactions; (ii) layouts of the model; (iii) annotations of elements and interactions. In this ecosystem, a curation platform should acquire biomedical knowledge from literature, standardise it to systems biology formalism, and provide stable, versioned annotations. The role of a curator is to create high-granularity, annotated building blocks (*facts*) that can be used in model building and referenced by the related layout. Such a modular ecosystem using curated, versioned and identifiable content requires better tool interoperability and collaboration between layout editors, model curators, annotators, and the research community in general (Fig. 2).

**Fig. 2.**
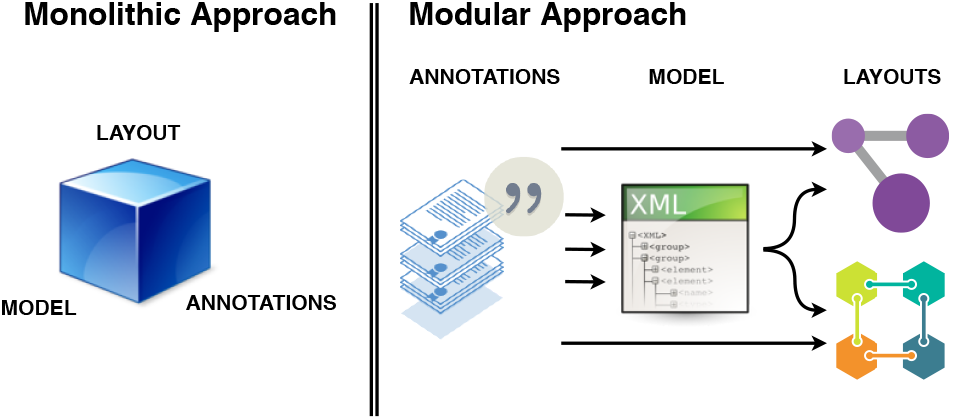
Monolithic hard building blocks hinder reuse and reference of content, while a modular workflow provides linkable content easier to reuse and share.

### A FACT AS A KNOWLEDGE UNIT

The concept of a *fact*, understood as a minimal piece of representative knowledge, can be found under various names in the literature. For example, Nano Publications [11] employs a fine-grained model, where a fact, called statement, consists of three basic elements: an assertion, its provenance and publication information. In BioNotate, facts are named *snippets*, defined as “*small chucks of text that may confirm or rule-out a relationship between two known entities*” [12].

Hence, a *fact* is a piece of knowledge that can be cited, referenced and attributed. In systems biology, a fact also needs to be serialised to a common format (e.g. SBML, RDF [13]) using ontologies for term normalisation in order to enable interoperable model building. Because a model of a biological process usually represents interactions of multiple components, such a model may consist of one or multiple *facts*.

### BIOKC WEB-BASED TOOL

In this paper we developed *BioKC* [14] (<u>Bio</u>logical <u>K</u>nowledge <u>C</u>uration) (Available at https://biokb.lcsb.uni.lu), a web-based collaborative tool for *fact* building, annotation and review. *BioKC* allows recording annotation and evidence sentences, storing them in a SBML-compliant data model, enabling the user to decide the granularity of their facts. *BioKC* provides a systematic workflow to ensure high quality control of the curation process. Once a fact reaches maturity, it can be released with stable uniform resource identifiers (URIs) and referenced from encoded layout or other tools.

*BioKC* is built on top of BioKB [15], a web-based interface designed to browse the text-mining results of almost 30 million publications including both abstracts and full-text articles. *BioKC* is a novel platform that enables the user to construct building blocks of systems biology models and allows the user to annotate them with human-provided and machine-identified literature evidence. Knowledge representation in *BioKC* follows the SBML standard in formalising elements of a given model, their relationships and annotations.

In the following Sections, we briefly review the current state of the art and approaches for model curation and annotation. Next, we describe the functionalities and design of *BioKC* supported by use cases. Finally, we describe further steps foreseen in the development of the platform.

## Related work

Literature on tools for annotation and curation of biomedical publications includes many platforms that, despite being described with similar keywords, showcase a broad range of features and purposes. In general, these tools are either suited for the annotation and curation of publications or focused on visual model building.

### ANNOTATION AND CURATION OF PUBLICATIONS

Different tools for knowledge curation were reviewed in two thorough surveys from Neves et al. following a detailed evaluation criteria. The first one [16] is specifically devoted to biomedical literature annotation tools, featuring 35 criteria used to evaluate 13 annotation tools [16]. Although this survey makes a distinction between text annotation (i.e. a complete tagging of a given text) and text curation (i.e. document analysis with respect to a given context), it does not consider model curation from the annotated/curated text information as a feature. Most of these tools are no longer available or do not feature collaborative web functionalities.

A recent survey from the same authors ([17]) covers a higher number of annotation tools but not all of them are directly related to the biomedical sciences [17]. In this case, 78 tools were selected following 26 criteria, and 15 tools were evaluated in detail. Most of the discarded tools were not available or were not web-based. Finally, only BioQRator [18], ezTag [19], MyMiner [20], tagtog [21], were suitable for biomedicine. However, none of these four tools support model curation or systems biology formats.

Table 2 summarises the criteria used in [17] except the publication impact of the tools, as these are not relevant for this work. Importantly, the criteria from Table 2 will be used for the description and evaluation of our tool, *BioKC*, in the Section <u>*Technical and Functional Comparison*</u>. We extended the table by including the tools relevant for curation in systems biology and biomedicine. In particular, we consider systems biology diagram editors and viewers as tools for curation because of their capability for model building and review.

**Table 1.**
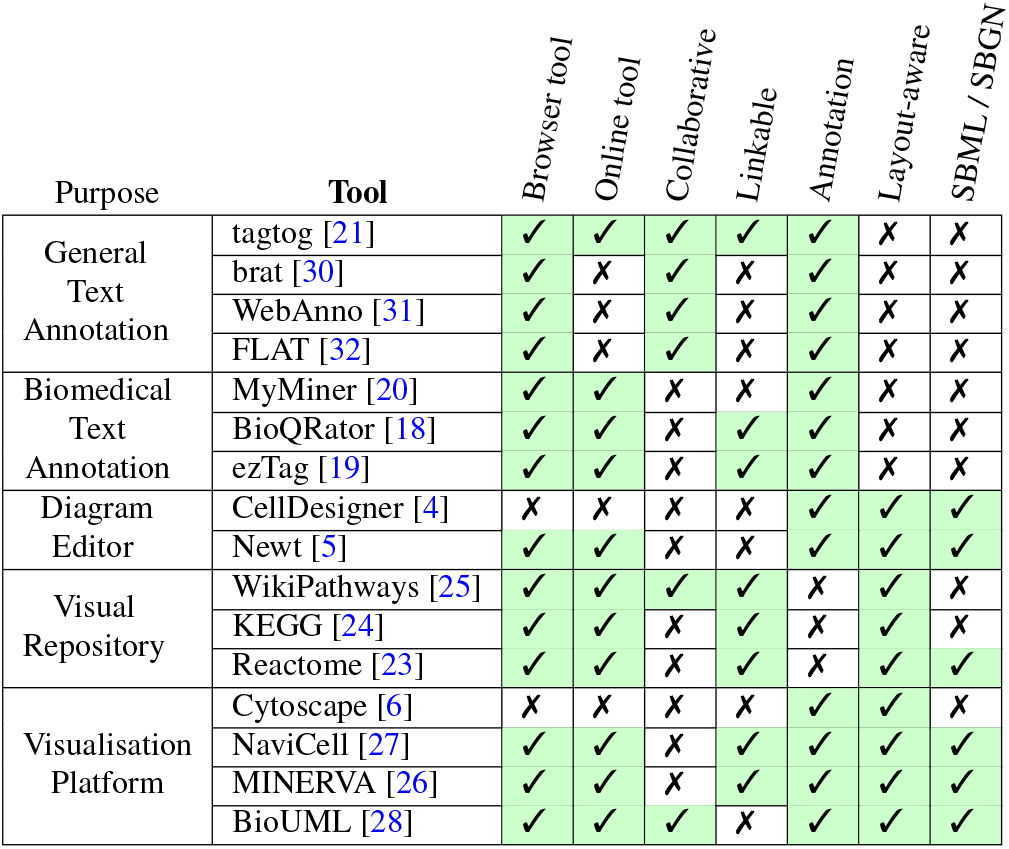
Summary depicting purpose, online availability and capabilities of different tools. Some browser-based tools are not available online, this distinction is shown in the first two columns. Collaborative column states which tools allow multi-user simultaneous operation. Linkable criterion refers to the ability to share and use the tool output as annotable content via URI-like links. Conversely, the annotation criterion indicates if a tool is able to produce annotations on the content.

**Table 2.**
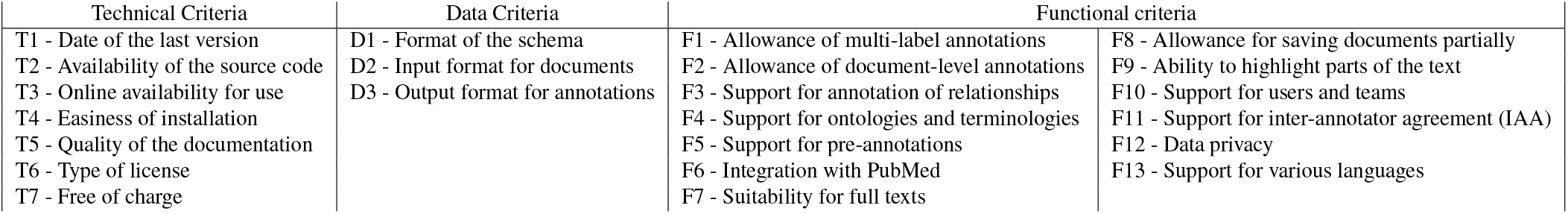
Technical, data-related and functional criteria) [17]. Publication criteria has been excluded as they do not apply for the comparison conducted in this paper.

### EDITING AND VISUALISATION OF MODELS

Many existing tools are able to create and parse systems biology diagrams encoded in different formats (e.g. SBGN, BioPAX) allowing the user to curate a layout or annotate a model. These tools are valuable in model curation, because the diagrammatic representation gives a comprehensive overview of the constructed model. The authors in [10] present a comparison of software tools suited to work with diagram layouts in systems biology standard formats [10]. They differentiate between **diagram editors**, like CellDe-signer [4, 22], Newt [5], or Cytoscape [6], and **management platforms** which include pathway databases as Reactome [23], KEGG [24], or WikiPathways [25]; and platforms for visualization of contextualised networks like MINERVA [26], NaviCell [27] or BioUML [28]. An important drawback of diagram-based model building is annotating the content. Diagram editors have limited capability to provide supporting evidence. Even though modelling languages support standardised annotations [29], they are difficult to introduce and maintain.

### SUMMARY

The ecosystem of tools for systems biology curation (see Table 1) offers solutions for publication annotation, layout editing, and knowledge exploration. Nevertheless, there is a lack of platforms for quality controlled model curation allowing online collaborative work. Some publication annotation tools like BioQRator support collaborative curation, but do not offer model building features. **Web repositories** and databases like BioModels [33] and CIDeR [34] host a multitude of models that can be downloaded in SBML, but these are *read-only* services. Interestingly, PathText2 [35, 36] is a step in the right direction, as it was designed to annotate biological pathway models with supporting knowledge from the literature, using SBML contents to query multiple databases and text-mining services.

In the light of the state of the art of curation tools for model building, the motivation for *BioKC* is threefold. First, we aim to provide a web application for collaborative model curation and annotation. Second, we want to implement features for a systematic curation workflow that will facilitate knowledge building and increase quality control. Finally, we seek to decouple the tasks of knowledge curation and of diagram building in systems biology. In this scenario, annotated and reviewed building blocks, *facts*, can be used as a linkable content to annotate model-based diagrams.

## Results

### FEATURES

#### Structure of a fact

In *BioKC*, a *fact* acts as an enriched SBML model composed of SBML elements such as species, compartments and reactions, as well as of other elements (annotations, cross-references) to support additional features. In particular, a *fact* belongs to a specific group of users, contains tasks and a change log that records actions. *BioKC* allows editing facts and the properties of their elements via web interface, adding more species, compartments or reactions (see Figure 3).

**Fig. 3.**
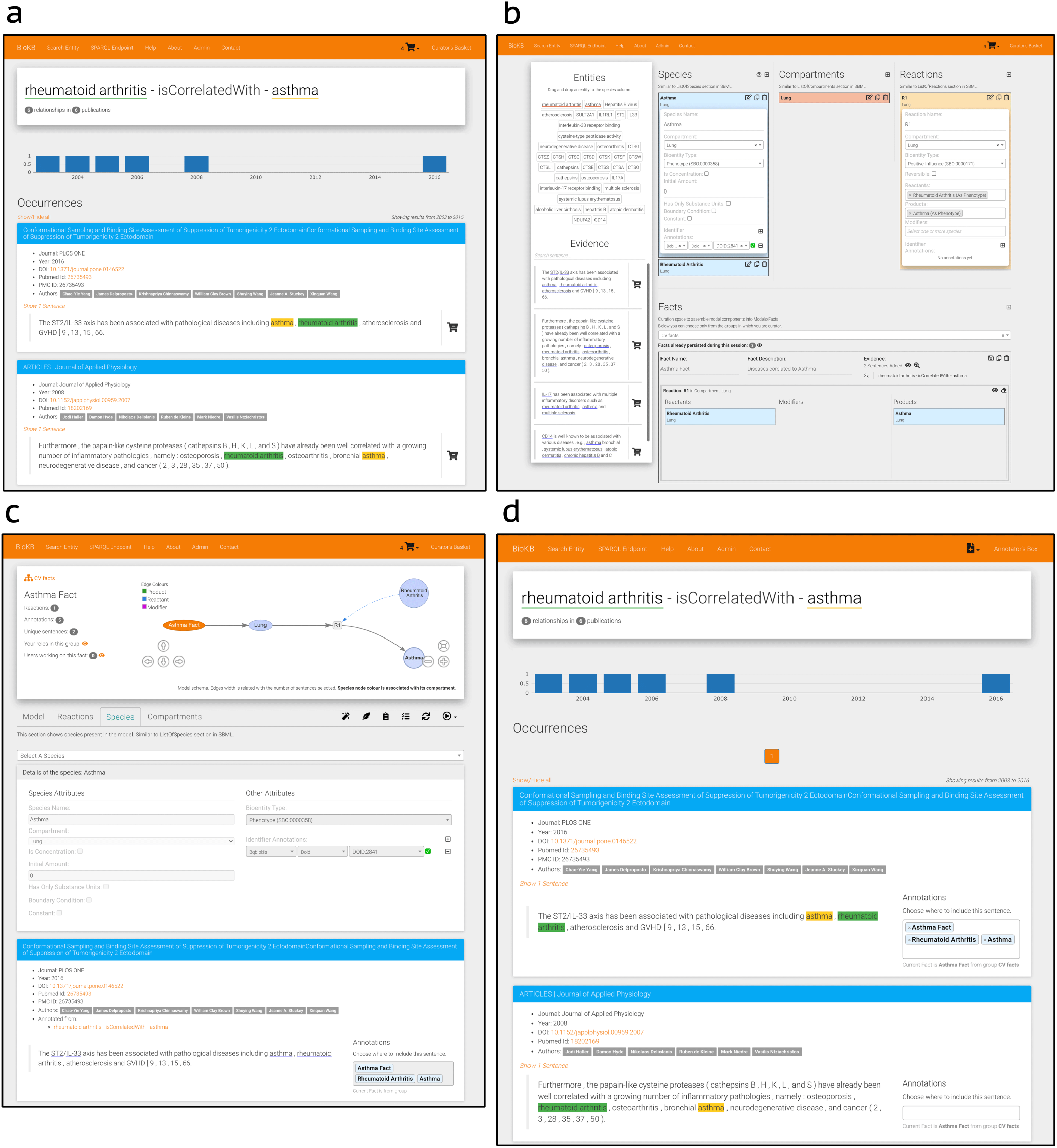
BioKC interface and functionalities. (**a**) BioKB relationship view showing sentences for a given entity relationship, sentences can be added to the basket. (**b**) Basket checkout redirects to the basket view where facts and their elements can be composed. (**c**) The fact view is where facts can be edited, either from scratch or after being persisted in the basket view. (**d**) The annotation mode enables annotation capabilities in BioKB to assign supporting evidence to one or multiple elements of a fact.

#### Annotation of a fact

All SBML elements inisde a *fact* can be further described, thanks to resolvable identifiers. *BioKC* supports the annotation of elements with BioModels qualifiers (https://co.mbine.org/standards/qualifiers) and Identifiers.org service [37], which includes more than 700 different namespaces. Such elements can also be annotated with supporting evidence from the literature either from BioKB or third party sources. Sentences from third party sources can be imported in both the basket and fact views (see Fig. 3 **b** and **c**). Basket mode supports bulk import of sentences from TSV files. Conversely, single sentences from third party sources can be added on the fact view. Both modes require certain provenance information. Namely, a valid DOI, PMC or PubMed ID must be provided to retrieve the corresponding publication metadata for the annotation.

#### Fact Groups and Multi-user Workflow

For a flexible management of the facts, these can be gathered in private groups. Users may be members of multiple groups and have different roles on each group. Group managers can grant read, annotation, curation, or management permissions to other members of the group. A warning will be shown if a user tries to curate/annotate a fact while another user is working on it.

#### Role system

Roles enable different sets of actions and are assigned per group of facts. Therefore, each user can have different set of roles for each group of facts. **Managers** can administrate their groups and member permissions as well as deleting facts or decide about the completeness of a task. **Curators** are able to add/edit/delete elements composing a fact (e.g. species or reactions). **Annotators** can assign sentences to elements of a fact. Finally, **readers** can inspect facts but cannot modify any aspect of them.

### USAGE

*BioKC* enables the user to work in two different directions: by first collecting the evidence, then creating facts from it (see blue box in Fig. 4); or, instead, starting with the curation of a fact and then annotating it with supporting evidence. Note the icon 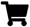 on the top right corner in Figure 3 **a** and **b**. This corner indicates the current operation mode.

**Fig. 4.**
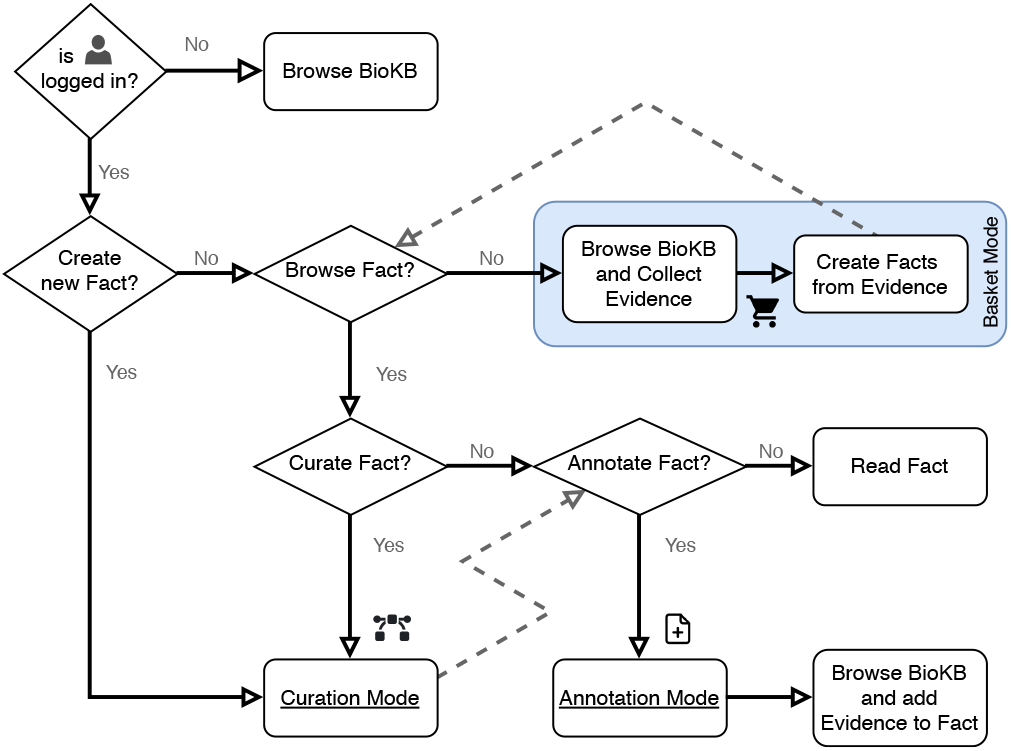
Flowchart describing the user operation flow and the different operation modes. The blue box shows the basket mode, which is the default operation mode when both curation and annotation modes are disabled.

The default mode is the *basket mode* for collecting evidence. Note how *BioKC* integrates with BioKB by adding the icon 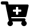 on the sentences (Fig. 3 **a**) enabling the user to collect evidence as it browses BioKB (or from third party source TSV file) and then compose multiple facts from the collected sentences in the *Basket view* (Fig. 3**b**). Once a fact is persisted, it can be further edited in the *Fact view* (Fig. 3**c**). Conversely, the second workflow entails the opposite procedure, which is depicted in the bottom images **c** and **d** of Figure 3. Lastly, *BioKC* also supports importing a model from a SBML file.

#### Curation

Users with curation permissions can start the *curation mode* from the fact view (see 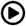 in Fig. 3 **c**) to add, delete or edit the elements that compose a fact. This mode also enables the use of aforementioned element annotations from resolvable identifiers. Top right corner will show the icon 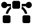 indicating curation mode is enabled.

#### Annotation

Similarly, annotators can start the *annotation mode* to add or remove supporting evidence from a fact. Such sentence annotations can be assigned to one or multiple parts of a fact, including the root element. Image **d** in Figure 3 shows how sentences in BioKB include a select box while the annotation mode is enabled. This mode also enables the annotator box (see icon 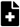 in Fig. 3 **d**) which lists recently visited pages. Sentences from external sources can be imported from the fact view using the *custom sentence annotation tool* (see 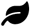 in Fig. 3 **c**).

#### Review

*BioKC* provides quality control and review mechanisms for the curation and annotation of facts. In particular, group managers can assign tasks to users from the *fact view* (see icon 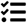 in Fig. 3 **c**) to guide the curation and annotation of the fact. Such tasks feature an annotator agreement system to assess the task completion. Users can also exchange messages and cast votes regarding their agreement or disagreement with the task completion. Managers have the final word over the task completion casting the *mark as finished* vote (see 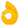 in Fig. 5). Once certain positive quorum is met, the task is marked as completed.

**Fig. 5.**
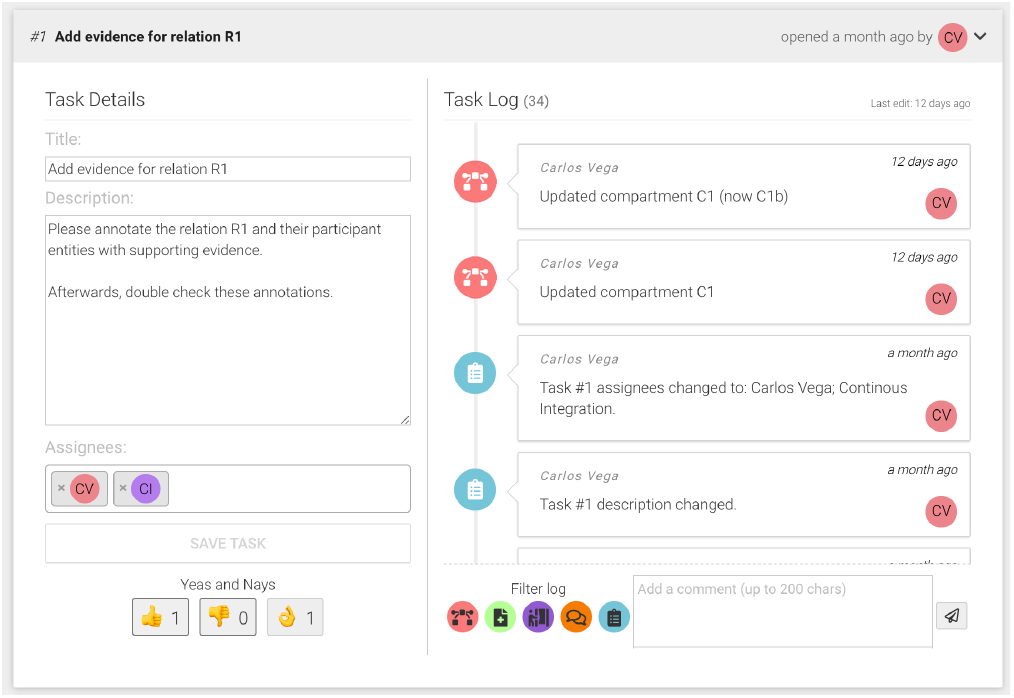
Example of task showing the title, description, assigned users and cast votes on the left. The right side shows the task log and comment input box.

## Discussion

High quality curation is key to provide reliable systems biology models. User-friendly annotation, collaborative features and quality control mechanisms are essential for such task. *BioKC* facilitates the process of curating annotated models in a standard and interoperable format as SBML.

Table 1 showcases the tools offering curation capabilities. However, although most are web-based, collaborative features are offered by only a few, mainly text annotation tools. *BioKC* was designed bearing in mind many capabilities from the diverse range of tools available, and particularly those closely related to text annotation tools. Below we present a detailed comparison of text annotation aspects.

### TECHNICAL AND FUNCTIONAL COMPARISON

Here, we use the criteria from [17] to compare technical, data-related and functional aspects of *BioKC* and other tools. In the original evaluation, points were assigned for completely (1), partially (0.5) or not (0) fulfilling a criterion. The sum of points was divided by the number of criteria, with a maximum score of 1. In the evaluation, tools obtained an average score of 0.62. Three best tools were WebAnno [31] (0.81), brat [30] (0.75) and FLAT [32] (0.71). Besides, a dedicated section was included for tools suitable for biomedicine, including ezTag (0.67), BioQRator (0.58), tagtog (0.6) and MyMiner (0.52).

We have included these 7 tools in our comparison and re-calculated the scores excluding the criteria not applicable for this paper: year of last publication (P1), citations in Google Scholar (P2), citations for corpus development (P3). Results can be found in Table 3, showing that *BioKC* coverage of the evaluation criteria is higher than other annotation tools, including those suitable for biomedicine.

**Table 3.**
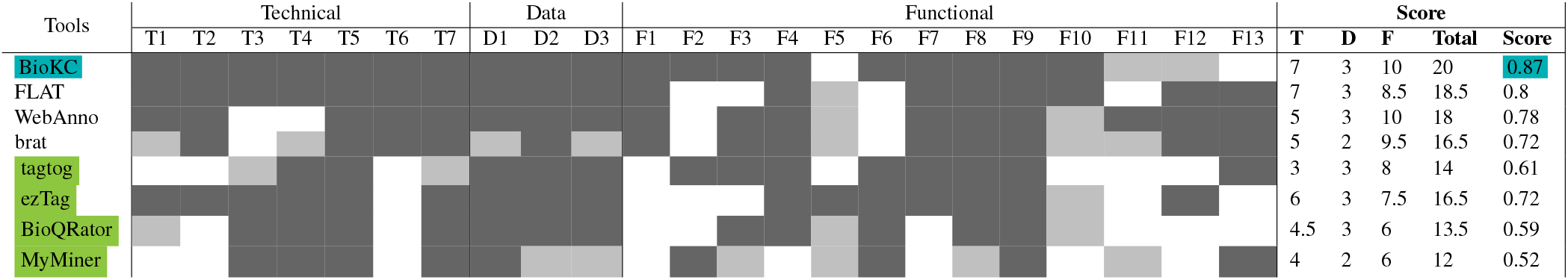
Criteria comparison of technical and functional aspects following criteria from Table 2. Gray colour indicates total fulfillment, light gray partial fulfillment, and white cells mean no fulfillment. Tools are sorted by descending score. Publication annotation tools suitable for biomedicine are highlighted in green.

Nevertheless, some criteria for *BioKC* are either partially fulfilled (F11, F12) or not fulfilled at all (F5, F13). Namely, F11 criterion is just partially satisfied, since, even though *BioKC* provides mechanisms to ensure certain level of inter annotator agreement over the curation and annotation process, it does not entail a fully blind annotation and curation workflow. Similarly, F12 criterion can be fulfilled as long as curated facts are kept private to their group members, but not once a fact is publicly released. F13 criterion is not satisfied since the platform dictionaries are in English. Lastly, F5 criterion is not met since annotation import is not supported yet. In summary, *BioKC* covers all technical and data criteria and most of the functional aspects of text annotation tools in [17].

### CURATION GUIDELINES COMPLIANCE

Notwithstanding, this evaluation compares tools regarding their annotation capabilities, while their main purpose differs from the aims of BioKC. Consequently, these criteria are not entirely exhaustive as some capabilities offered by *BioKC* are not covered. Such capabilities have been described separately in previous sections (see Section Features).

To complete the assessment of *BioKC* we made use of on a recent work from [38] introducing the Minimum Information about a Molecular Interaction CAusal STatement (MI2CAST). MI2CAST consists of rules and good practices for the curation of causal molecular interactions. The first three rules cover mandatory information about the interaction: i) the source and target entities, ii) the effect of the interaction, and iii) the evidence provenance. Additionally, the fourth rule recommends encoding contextual information. MI2CAST guidelines do not impose a particular format in which interactions should be represented or encoded. Therefore, we strongly believe that features and capabilities of *BioKC* described in this paper comply with MI2CAST guidelines and recommendations.

### CONCLUSIONS

We present *BioKC*, a web-based platform for collaborative curation and annotation to cope with the new needs of systems biology models building. Our platform offers quality control and reviewing features for curation and annotation that are not available in the current state of the art. *BioKC* platform is in constant development and its roadmap (https://biokc.pages.uni.lu/roadmap/) foresees support for defining and annotation of complexes, and handling of SBML extensions such as the Multistate and Multicomponent species package [39]. Supporting a wider range of text-mining knowledge bases and model repositories is essential for *BioKC*’s commitment with interoperability. With our work we aim to ease research collaboration providing features to review the curation process and quality control the annotation of supporting evidence. Thus, eventually reducing the time and effort needed to build the models.

## Materials and Methods

### ARCHITECTURE

The proposed solution, *BioKC*, extends BioKB functionality. BioKB is a platform designed to help researchers easily access semantic content of millions of abstracts and full text articles [15]. BioKB features a text-mining pipeline that extracts relations between a wide variety of concepts, including proteins, chemicals, diseases, biological processes and molecular functions, providing directionality when possible to infer. Extracted knowledge is stored in a knowledge base publicly accessible via a web interface and a SPARQL endpoint. Figure 6 summarises BioKB and *BioKC* workflow for model curation with disease maps as a use case example of such curation workflow. As a result, *BioKC* allows the use of both machine extracted knowledge from BioKB and human-provided annotations, enabling the user to introduce annotations from external sources. Below we describe the design of *BioKC*’s features.

**Fig. 6.**
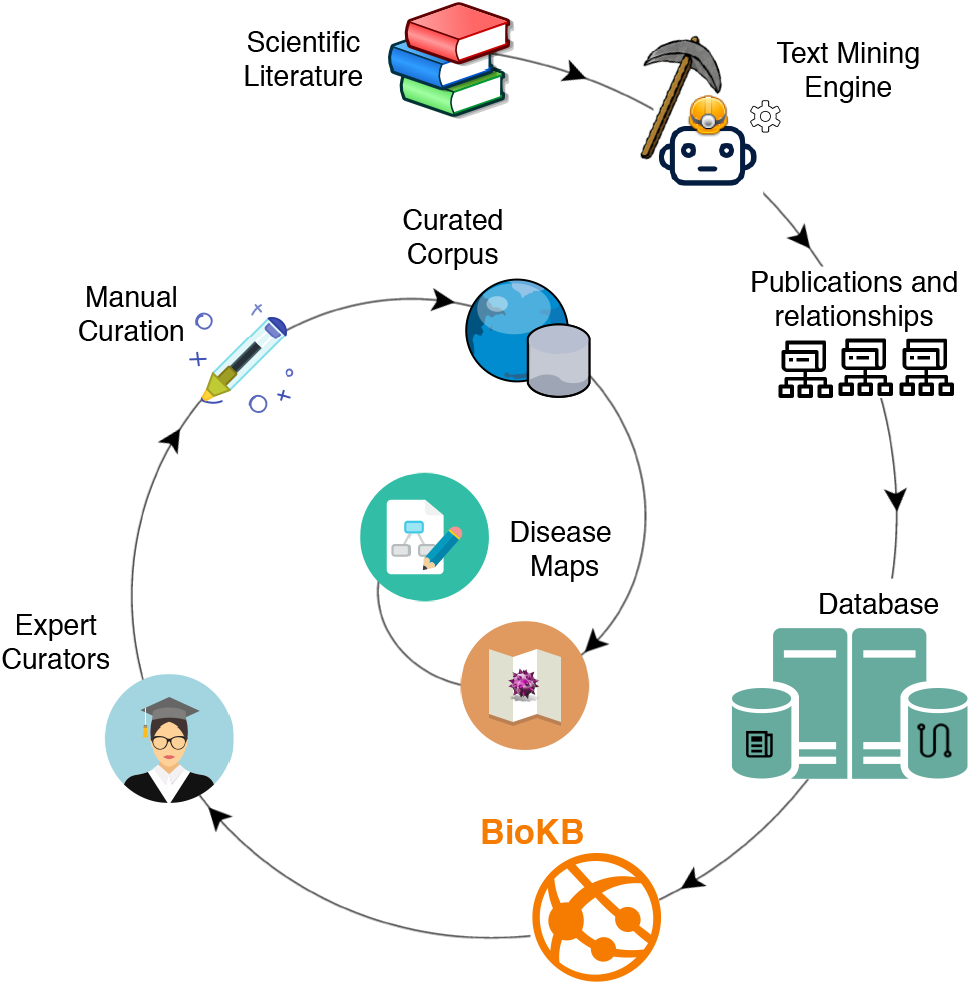
BioKB and *BioKC* pipeline for the curation of systems biology models. An example use case of *BioKC* is the development of disease maps.

#### Implementation Environment

*BioKC* is developed in Python and JavaScript, which allow for fast iterative development life cycle in both front-end and back-end. Flask and SQLalchemy are employed for the web server and database implementations, with Vue, Jquery and other JavaScript libraries contributing to a real-time collaborative and interactive multi-user experience in the client side.

#### Multi-level annotation

*BioKC* follows a SBML-like data model in which every object composing a SBML model inherits all properties from SBase abstract type depicted in Figure 7. This hierarchy was replicated using SQL joined table inheritance polymorphism. Hence model tables like Species, Compartment, Reaction, SpeciesReference, etc. inherit these properties allowing the model to be annotated at different levels (i.e. annotations can be assigned to compartment, species, etc.).

**Fig. 7.**
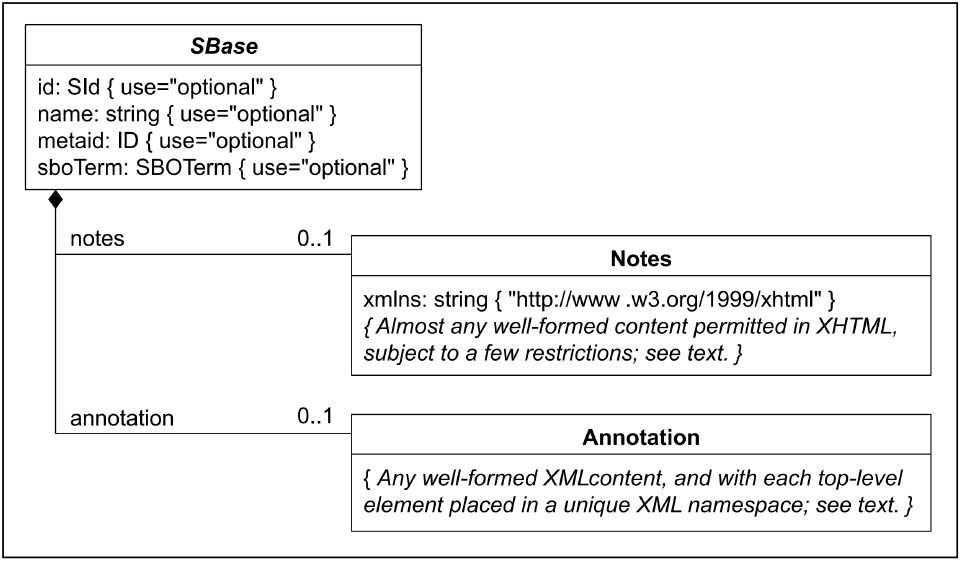
Nearly every object composing an SBML Level 3 model definition has a specific data type that is derived directly or indirectly from a single abstract type called SBase. See Section 3.2 from SBML Specification for Level 3 Version 2 Core. *BioKC* follows the same structure for all SBML elements composing a *fact* so that they can be annotated.

#### Multi-granular Fact Curation

The concept of *fact* (see Section <u>*A Fact as a Knowledge Unit*</u>) is employed in other tools as a way to encode pieces of knowledge. Nonetheless, *BioKC* makes use of a known standard format like SBML, as it provides versatility on the model size, allowing to encode very large models, or small pieces of curated knowledge. *BioKC* provides similar versatility by following above-mentioned polymorphic design, letting the user decide the structure and extent of their facts.

#### Action log

Multi-user collaborative work requires registering the actions undertaken on the model. For this, we employ a custom declarative base class and a SQLalchemy *mixin*, allowing adding common columns to multiple tables that share this functionality. Specifically, each table has two columns `created_on` and `updated_on`, that register the creation date and last modification time, respectively. Benefiting from previously described data model polymorphism, each action is registered in the modified element as a Note (see Fig. 7) with a User as author and a comment describing the action. Such actions can be assigned to multiple tasks to better organise the actions taken during the curation process.

## ACKNOWLEDGEMENTS

The authors would like to thank the Luxembourg National Research Fund (FNR) for supporting this work through grant 14729738 for Covid19 Literature Biocuration, Text-mining And Semantic Web Technologies (COVlit) and the National Centre of Excellence in Research on Parkinson’s disease (NCER-PD [FNR/NCER13/BM/11264123])

## AUTHOR CONTRIBUTIONS

**Carlos Vega**: Conceived and designed the development roadmap; developed the solution; wrote the manuscript.

**Valentin Grouès**: Conceived and designed the development roadmap; developed the solution; contributed and reviewed the manuscript.

**Marek Ostaszewski**: Conceived and designed the development roadmap; supervised the work; contributed and reviewed the manuscript.

**Reinhard Schneider**: Supervised the work; contributed and reviewed the manuscript.

**Venkata Satagopam**: Conceived and designed the development roadmap; supervised the work; contributed and reviewed the manuscript.

